# Wheat Panache - a pangenome graph database representing presence/absence variation across 16 bread wheat genomes

**DOI:** 10.1101/2022.02.23.481560

**Authors:** Philipp E. Bayer, Jakob Petereit, Éloi Durant, Cécile Monat, Mathieu Rouard, Haifei Hu, Brett Chapman, Chengdao Li, Shifeng Cheng, Jacqueline Batley, David Edwards

## Abstract

Bread wheat is one of humanity’s most important staple crops, characterized by a large and complex genome with a high level of gene presence/absence variation between cultivars, hampering genomic approaches for crop improvement. With the growing global population and the increasing impact of climate change on crop yield, there is an urgent need to apply genomic approaches to accelerate wheat breeding. With recent advances in DNA sequencing technology, a growing number of high-quality reference genomes are becoming available, reflecting the genetic content of a diverse range of cultivars. However, information on the presence or absence of genomic regions has been hard to visualize and interrogate due to the size of these genomes and the lack of suitable bioinformatics tools. To address this limitation, we have produced a wheat pangenome graph maintained within an online database to facilitate interrogation and comparison of wheat cultivar genomes. The database allows users to visualize regions of the pangenome to assess presence/absence variation between bread wheat genomes.

Database URL: http://www.appliedbioinformatics.com.au/wheat_panache

## Introduction

Bread wheat (*Triticum aestivum*) is one of the most widely grown crops, yet there is a significant challenge to increase yield to meet the projected demands of a growing world population. With predictions of climate change-related yield losses ranging from 17% to 31% by the middle of the 21^st^ century (1), improved genomics-based breeding approaches are required to produce climate change-ready wheat cultivars.

Wheat genomics has made rapid advances in recent years with the first draft genome assembly produced in 2014 (2) based on the shotgun sequencing of isolated chromosome arms (3–5). A first near-complete assembly of the variety Chinese Spring was produced in 2017 (6) with a final reference genome assembly available in 2018 (7). This reference assembly was rapidly followed by assemblies of fifteen additional cultivars from global breeding programs (8).

The increasing availability of reference genome assemblies made it clear that there is significant presence/absence variation (PAV) between individuals (9–12). This insight has led to the production of pangenomes that reflect the gene content of a species rather than an individual (10,13–20). Pangenomes are now available for several plant species; the first bread what pangenome representing the gene content of 16 bread wheat cultivars was published in 2017 (18). This wheat pangenome was assembled using an iterative mapping approach, which efficiently identified new gene space and called gene presence/absence between individuals. This kind of pangenome is however limited in that the physical location of the new gene space can be difficult to determine with accuracy. With the availability of multiple whole-genome references, this limitation may be addressed through the production of a graph-based pangenome. Graph-based pangenomes have recently become popular thanks to the graph data structure which can accurately represent the physical locations of genomic and structural variants with minimal reference bias, with tools such as vg (21), seqwish (22), minigraph (23), and PHG (24) being successfully applied to build variation, sequence, or haplotype graphs.

A major limitation of pangenome graphs is that few tools are available to visualize these complex graph structures. Genome visualization tools such as GBrowse (25), JBrowse2 (26), or Circos (27) are designed to display information relative to a linear reference genome, not a graph of several genomes, while graph viewers such as Bandage (28) or pangenome viewers such as ODGI (29) focus on visualizing the graph itself, but display little other information such as genome annotations.

Panache is a recent pangenome visualization tool that can process linearized assembly graphs and display shared regions as a web-based dynamic heatmap (30). Panache has so far only been applied to visualize presence/absence variation in the banana pangenome (14), but has the potential to be expanded to other species, even for crop genomes as large as wheat. Here, we present a graph pangenome representing 16 bread wheat cultivars hosted within a public Wheat Panache database, with a new web-based browser for visualizing genomic regions across the wheat pangenome, along with the graph formatted for minimap2 (31) and Giraffe (32). This tool offers researchers and breeders the ability to assess genome variation between these varieties, mining the diversity present in this large and complex genome.

## Results and discussion

### A wheat graph pangenome

We constructed a graph pangenome using 16 high-quality wheat genome assemblies Representing the global variation of modern bread wheat cultivars. The assembled graph had a total size of 15.8 Gbp in comparison with the founder genome assembly sizes of 13.9 to 14.2 Gbp (33). After aligning all genomes back to the graph, these 15.8 Gbp were split up into 2,791,482 segments present in at least one individual. The segments had an average size of 5.6 Mbp (median: 498 bp), ranging from 2 bp to 37.6 Mbp (Figure 1A). Realignment of the 16 genome assemblies to the graph revealed that out of the 2.7 million segments, 542,711 (19%) segments were present in all individuals (total size: 10.2 Gbp ranging from 2 bp to 4.9 Mbp) with the remaining 2,248,771 segments (total size: 5.6 Gbp) being present in a median of 8 individuals (Supplementary Figure 1). 10,437 segments (0.4% of all segments) with a total length of 19.9 Mbp (average length: 1.9 Kbp) were not covered by any genome assembly during the realignment step, probably due to these segments being too small and/or too repetitive.

**Figure 1:**
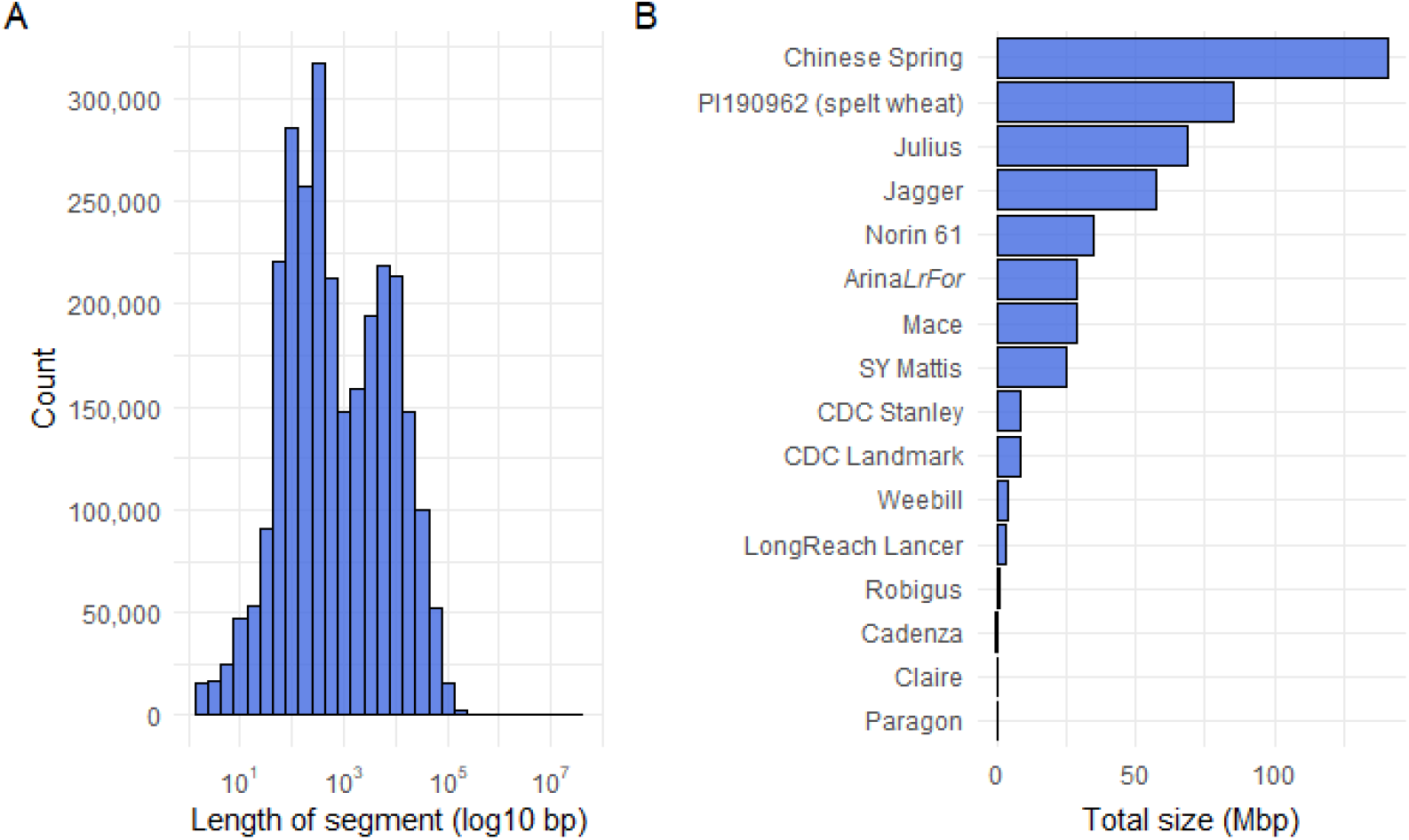
A) Bar chart showing the distribution of the size of all assembly graph segments (log-scale). B) Total size of unique segments per cultivar in Mbp. PI190962 is a line of species *Triticum spelta*, Chinese Spring is the reference cultivar of *T. aestivum*.

Interestingly, the cultivar with the most unique segments was the reference cultivar Chinese Spring, with 158,503 (7%) of segments with a total size of 140.5 Mbp being only present in Chinese Spring (Figure 1B). This may be due to the genomic distance between Chinese Spring and the other cultivars, consistent with previous observations (18), and reflecting Chinese Spring’s age (collected around 1900) and its lack of agronomic characters that were selected for in modern cultivars (34). The distance between the Chinese Spring assembly and the 15 other assemblies is also supported by 1.2 Gbp of the graph in 901,475 segments not being present in Chinese Spring but in at least one other cultivar, reflecting the complex history of introgressions in modern bread wheat (8,35). We aligned the IWGSC v1 gene annotation for Chinese Spring (7) back to the graph by intersecting the linearized graph with gene positions. We found a position in the graph for 110,790 (100%) genes confirming that the graph assembly contains all gene models of the IWGSC assembly.

### The Wheat Panache web portal

Using this graph, we built a web-based Panache instance (30), allowing users to visualize regions or genes of interest for presence/absence across the chosen wheat cultivars. The webserver is available at http://www.appliedbioinformatics.com.au/wheat_panache.

Wheat Panache displays a linear version of the pangenome graph subdivided into blocks based on presence/absence of the selected individuals. A block is defined to have no internal presence/absence variation and to contain at least one gene. Blocks are named based on the pseudomolecule they originated in, and as we started the assembly with the IWGSC assembly, most blocks (1,890,035 out of 2,791,483 blocks, 67%) are named after their position in the IWGSC assembly.

The interface displays the linearized pangenome as a chain of such graph segments, with one horizontal track per cultivar (Figure 2). Coordinates are based on the pangenome graph assembly. Genes are represented as black dots above blocks and hovering over a gene reveals its coordinates within the assembly and exon structure. Three summary tracks below the cultivar tracks show which blocks are core or variable based on a user-definable threshold, how long the block is, and how often the block is repeated within Panache. Users can zoom into blocks, or search for ‘hollow areas’ (areas of consecutive absence based on a user-defined threshold) using the Hollow Area Finder, which is a convenient way to automatically focus on large PAV areas. Users can sort the cultivars alphanumerically, by gene presence/absence status, or by a phylogeny based on Mash v2.3 (36). The graph assembly displayed in Wheat Panache, including a version pre-indexed for vg v1.37.0’s Giraffe (32) is available at https://doi.org/10.5281/zenodo.6085239 (37) allowing for downstream analyses of the population graph.

**Figure 2:**
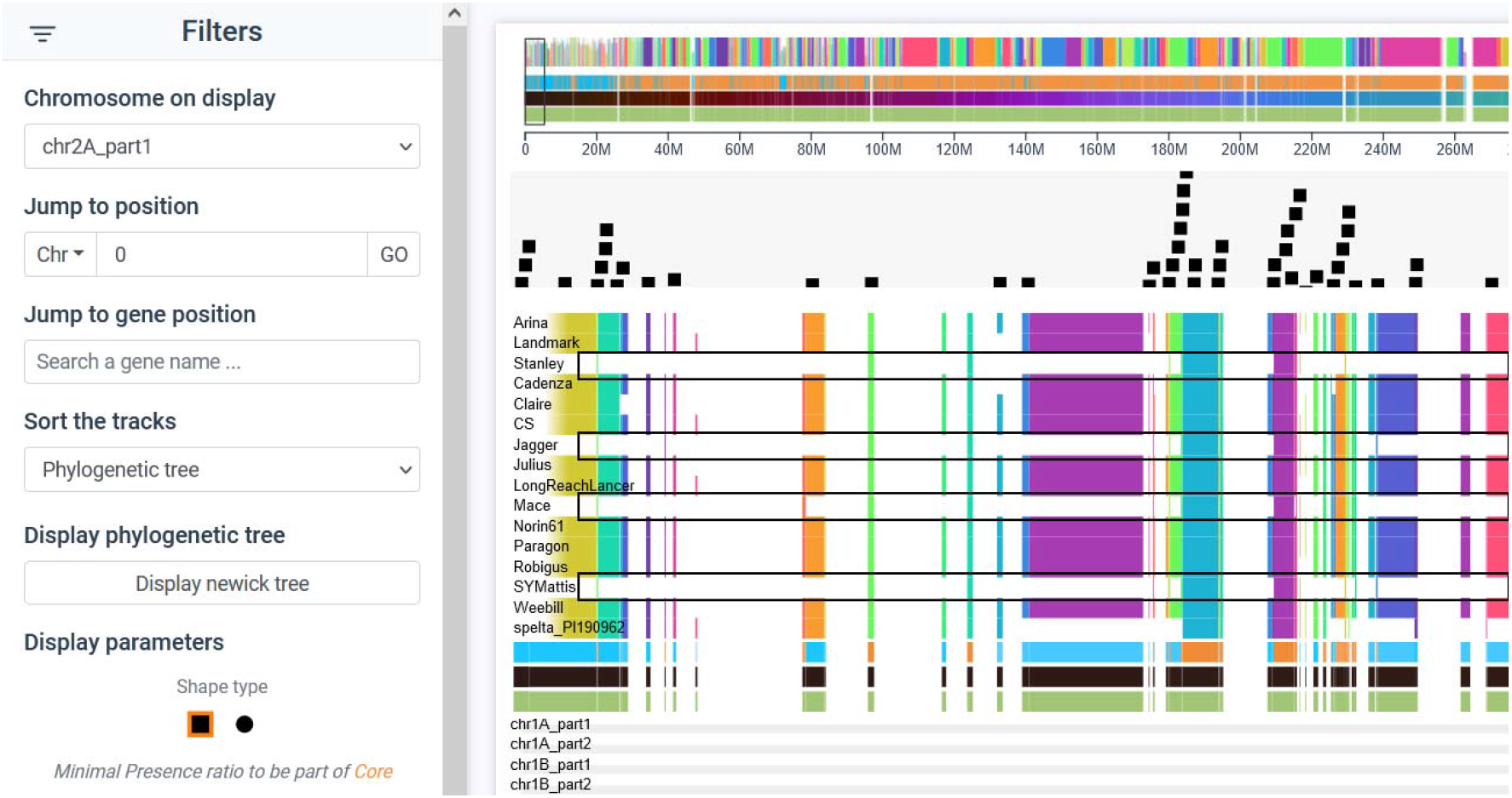
Wheat Panache screenshot showing a *Aegilops ventricosa* introgression at the beginning of chromosome 2 in cultivars Stanley, Jagger, Mace, and SY Mattis (39,40). Black boxes were added to show the region missing in cultivars where the introgression replaced parts of 2A. The graph assembly started with the IWGSC v1 assembly leading to linearized regions following the same naming scheme as the IWGSC v1.0 assembly (chr1A_part1, chr1A_part2, chr2A_part1, …). CS stands for Chinese Spring. Shown here is the beginning of the first part of chr2A. Black blocks are gene models. White regions correspond to regions that are present in the graph but contain no genes.

In summary, we present the first wheat graph pangenome assembly, based on 16 cultivars with an online visual representation of the graph within the Panache visualization tool. The graph assembly will be a valuable tool for wheat genomics researchers looking for a more accurate reference assembly. The web platform Panache allows users to interrogate this graph and search for structural variants around regions of interest.

## Materials and Methods

We used publicly available genome assemblies, including fifteen high-quality *Triticum aestivum* genome assemblies (8) and the IWGSC v1 *T. aestivum* cv. Chinese Spring assembly (7), to assemble a graph using minigraph v0.14 (23). To optimize assembly, we used k-mers that appear fewer than 100 times (-f.1) for the graph assembly and assembled the graph genome by genome, starting with IWGSC v1 followed by alphabetical order of cultivar names, ending with the *T. aestivum ssp. spelta* PI190962 assembly.

All assemblies were aligned with the final graph using minimap2 v2.18 (31) and alignments were converted to BED format. The main graph was linearized using gfatools gfa2bed v0.4 with default parameters (https://github.com/lh3/gfatools/releases) and merged with all minimap2 alignments using bedtools v2.30.0 multiinter (38). The resulting blocks were intersected with the IWGSC gene annotation using bedtools v2.30.0 intersect.

The data was converted to Panache JSON format and a Panache instance was set up to serve the data (30). To make the display feasible on a regular workstation, we retained only blocks overlapping with Chinese Spring genes and then merged adjacent blocks if they showed identical PAV behaviour across all individuals.

## Acknowledgments

This work is funded by the Australia Research Council (Projects DP210100296, DP200100762, and DE210100398). This work was supported by resources provided by the Pawsey Supercomputing Centre with funding from the Australian Government and the Government of Western Australia.

